## Supplementary Material for "Targeted V1 comodulation supports task-adaptive sensory decisions"

### Targeted V1 comodulation supports task-adaptive sensory decisions Supplementary Material

#### S1 Statistics of V1 population responses

We summarize single unit statistics across the V1 population and look for differences between task-informative neurons and task-uninformative neurons. We find that the distributions of mean firing rate in a trial is shifted slightly higher for the informative neurons (Fig. S1A). This is expected, since activity is necessary to find systematic activity differences with stimulus orientation. Informative units have an average rate in a trial of 42 spikes/second with a standard deviation of 22, while uninformative units have a mean of 37 spikes/second with a standard deviation of 20. We also find that the distributions of Fano factors for high contrast stimuli is shifted slightly higher for informative neurons (Fig. S1B, informative units have a mean Fano factor of $= 1.14 +/- 0.47$, uninformative units have a mean Fano factor $= 1.06 +/- 0.62$).

#### S2 PCA is insufficient to robustly find targeted modulation

We ask whether a simpler analysis based on dimensionality reduction (principal component analysis, PCA) of the fitted SR model residuals would suffice for robust modulator estimation in the V1 data. We find that neither the eigenspectrum of the data (Fig. S2A), nor that of the residuals (Fig. S2B) reveal low-dimensional structure. Nonetheless, the first principal component roughly aligns with the projection into latent space, $\mathbf{C}$ estimated by the modulated SR model (Fig. S2C) providing a noisy version of the estimated modulator coupling (Fig. S2C). We further find that PCA results are highly dependent on the stimulus condition. Depending on whether PCA residuals are computed on only high contrast stimulus residuals or on any contrast stimulus residuals, the resulting PC axis can vary substantially (Fig. S2D, residuals computed from the SR model). Interestingly we also see that the variance explained by the first PC axis is smaller in the control task vs the relevant task if any stimuli are included, but larger if only high contrast stimuli are included (Fig. S2E-F). This suggests that standard dimensionality reduction of residuals might not be sensitive enough to detect low dimensional modulator structure. Taking the square-root of the activity before applying PCA (a common preprocessing step for homogenizing the data variability [1]), does not change the results qualitatively. Instead explicit latent dynamical models jointly fitted with a SR model are required to detect the modulator. This suggests that such low-dimensional targeted modulation may be more ubiquitous than one would expect from previously reported analyses.

#### S3 V1 modulation

**Modulator statistics** We tested whether the extracted V1 modulator could be exclusively caused by stimulus variations that are not sufficiently captured by our linear stimulus-response model. For this we look for variations in the statistics of the modulator with different stimuli. We computed the mean value of the modulator over all stimulus presentations in a block for low and high contrast stimuli and find that the distribution of modulator values is similar for different contrast conditions (Fig. S2G). We then compute modulator mean and variance respectively and separately for each of the four time windows of 50ms for which the stimulus was present. The mean of the modulator does not fluctuate much within stimulus presentations (Fig. S2H, top). Moreover, the differences between high and low contrast are small. The same applies to the estimated modulator variance which is similar within different time bins of the stimulus presentations and across the two contrast conditions (Fig. S2H, bottom). Overall, this suggests that the modulator is unlikely to be merely accounting for residual stimulus responses that were not captured by the SR model, but is instead consistent with a stationary fluctuating modulatory source that is independent of the stimulus.

### S4 Comparison to On/Off states

The modulator extracted from our data has the form of normally distributed noise (see Fig. S2G and Fig. S2I). Other types of modulation have previously been described in the literature [2], with different statistics. Specifically On/Off states modulate population activity in a binary manner, in the context of attention. We verify that our analysis is able to differentiate between these two different forms of modulation. Since the PLDS framework used for the modulated SR model assumes a Gaussian prior noise distribution (see details in Methods of main text), it is conceivable that the unimodality of the extracted modulator may be a consequence of this prior. In order to understand whether the influence of the prior could be strong enough to conceal true binary modulation, we fit the PLDS model to simulated data that has a binary ground truth modulator. We match the simulation statistics to our recordings, by using parameters estimated in an example session, but change the modulator statistics to reflect two discrete states. We simulate data from the resulting model and then repeat the same fitting procedure as that used on the real data. We find that the estimated modulation is strongly bimodal and reflects the binary modulator well, despite the model's prior (see Fig. S2I). We further find that the spiking patterns are qualitatively different in the simulations with the binary modulator, than in the real data, as they visually reflect the two different spiking regimes (see Fig. S2J). This does not exclude the presence of fast On/Off dynamics in the data but it suggests that the modulator here is distinct.

### S5 Partial correlation analysis for mean rate, coupling and informativeness

We perform a partial correlation analysis to account for effects of mean firing rate on the relationship between coupling and informativeness. A linear regression lets us average out the linear effects of mean firing on informativeness (Fig. S2K) which explains most of the relationship between the variables (Fig. S2L). When then correlating the residual informativeness (not explained by mean firing rate differences) with the coupling we find that the relationship is preserved (Fig. S2M-N).

### S6 Excess correlations

The modulated SR model captures part of the shared variability between neurons. To test for additional structure, we look at the pairwise correlations expected from the modulated SR model versus those in the data. We find that the pairwise correlations are slightly larger in the real data than what would be expected due to differences in the response statistics (when simulated from the modulated SR model). More specifically, we find that those excess correlations are, on average, slightly larger between pairs of informative units than pairs of uninformative or informative-uninformative units (Fig. S2O-R). This suggests that there is additional structure, not captured by the modulator, which may be related to other sources of noise correlations [3; 4].

### S7 Response modulation due to attention

It has previously been reported that neurons with receptive fields overlapping an attended stimulus increase their activity[5; 6; 7]. However, here we do not find evidence that the increase in activity due to attention or task condition is specific to the task-informative units. Specifically, the correlation between attentional modulation and task-informativeness is close to 0 in the populations we were able to compare (sufficient trials and sufficient number of informative neurons, see Methods for exclusion criteria) (Fig. S3A). The attentional modulation in a single neuron is measured as the difference between response to high contrast stimuli in the relevant task condition (attend-in) minus in the control task condition (attend-out) divided by the sum of the two [8]. Similarly there is no significant correlation in the modulator coupling strength and the attention index in any of the blocks we were able to compare (3 pairs of blocks in the same session with good modulator fit; Fig. S3B).

### S8 Adaptation

Within a trial, stimuli are flashed on (for 200ms) and off (for 200-400ms) at low or high contrast several times before the target appears. We exclude the first stimulus presentation from our analysis, to avoid effects of initial transients or adaption [9]. Nonetheless, we wanted to test whether adaptation could interfere with our estimates of the informativeness of a unit or the modulated SR model fits. For this we define a summary statistic for adaptation in single units as follows: we group the last repeat stimulus responses from all trials by the number of repeats that preceded them. We compute the average of each group and the sign of the differences between them. We average over the signs to obtain a value between -1 (decreasing in response with increasing number of repeats) and 1 (increase in response with increasing number of repeats). Fig. S3C shows the distribution of adaptation indices over all blocks and units, compared to that of the subset of blocks well fitted by the modulated SR model. This distribution is broad overall, with no significant differences between populations that are well fitted by a model including modulation versus units that are badly fit by the model. Similarly, we see no systematic relationship between the adaptation index and the informativeness of a unit (Fig. S3D). Overall, these results suggest that classic adaptation cannot trivially explain the effects of modulation or its targeting towards informative neurons.

### S9 Multiunits

Many of the responses in the analyzed dataset are likely a composite of multiple neurons, and we wanted to examine the effects this could have on our results. We use mathematical analyses and numerical simulations, both based on a simple encoding model (S3E), to ask how estimates of informativeness and targeted modulation are affected by pooling together the activity of several (potentially similarly tuned) neurons.

**Estimating informativeness from multiunits.** We model multiunit activity as the sum of activities of pairs of neurons, $i$,$j$, each described as a Poisson processes with spike count distribution Poisson $\left(\lambda_{i,j|s}\exp(c_{i,j}m_t)\right)$, as in the main text theory (Eq. 1). The informativeness of the multiunit may be written as:

$$d' = \frac{\left(\mu_{i|s_1}+\mu_{j|s_1}\right)-\left(\mu_{i|s_2}+\mu_{j|s_2}\right)}{\sqrt{\frac{1}{2}\left(\left(\sigma^2_{i|s_1}+\sigma^2_{j|s_1}+2\mathrm{Cov}_{i,j|s_1}\right)+\left(\sigma^2_{i|s_2}+\sigma^2_{j|s_2}+2\mathrm{Cov}_{i,j|s_2}\right)\right)}}, \tag{1}$$

where we have used the standard relationship for the variance of a sum of correlated variables.

Given the doubly stochastic nature of the single neuron responses (formally, a log-gaussian Cox process [10]), the average firing rate of neuron $i$ in response to a stimulus $s$ is:

$$\mu_{i|s} = \mathbb{E}\left[\lambda_{i|s}\exp(c_i m_t)\right] = \lambda_{i|s}\exp\left(\frac{c_i^2\sigma_m^2}{2}\right), \tag{2}$$

where $\sigma_m^2$ is the variance of the modulator. The corresponding variance is

$$\begin{aligned}\sigma^2_{i|s} &= \mathbb{E}\left[\lambda_{i|s}\exp(c_i m_t)+\lambda^2_{i|s}\exp(2c_i m_t)\right]-\mu^2_{i|s}\\ &= \lambda_{i|s}\exp\left(\frac{c_i^2\sigma_m^2}{2}\right)+\lambda^2_{i|2}\exp\left(2c_n^2\sigma_m^2\right)-\lambda^2_{i|s}\exp\left(c_i^2\sigma_m^2\right).\end{aligned} \tag{3}$$

Finally, the covariance of responses for neurons $i$ and $j$ given a stimulus $s$ is

$$\begin{aligned}\mathrm{Cov}_{i,j|s} &= \mathbb{E}\left[\lambda_{i|s}\lambda_{j|s}\exp\left((c_i+c_j)m_t\right)\right]-\mu_{i|s}\mu_{j|s}\\ &= \lambda_i\lambda_j\exp\left(\frac{(c_i^2+c_j^2)\sigma^2}{2}\right)\left(\exp(c_i c_j\sigma^2)-1\right).\end{aligned} \tag{4}$$

In the limit when the modulator variance is very small ($\sigma_m^2 \to 0$), we have $\mu_{i|s} \to \lambda_{i|s}$, $\sigma^2_{i|s} \to \lambda_{i|s}$ and $\mathrm{Cov}_{i,j|s} \to 0$, so the informativeness becomes:

$$d' = \frac{\lambda_{i|s_1}+\lambda_{j|s_1}-\lambda_{i|s_2}-\lambda_{j|s_2}}{\sqrt{\frac{1}{2}\left(\lambda_{i|s_1}+\lambda_{j|s_1}+\lambda_{i|s_2}+\lambda_{j|s_2}\right)}}. \tag{5}$$

We will now show that, in this limit, and assuming the neurons in the multiunit have the same stimulus preference, the absolute value of the multiunit informativeness is bounded from above by the sum of that of the two component neurons, i.e.

$$\frac{|\lambda_{i|s_1}+\lambda_{j|s_1}-\lambda_{i|s_2}-\lambda_{j|s_2}|}{\sqrt{\frac{1}{2}\left(\lambda_{i|s_1}+\lambda_{j|s_1}+\lambda_{i|s_2}+\lambda_{j|s_2}\right)}} \leq \frac{|\lambda_{i|s_1}-\lambda_{i|s_2}|}{\sqrt{\frac{1}{2}\left(\lambda_{i|s_1}+\lambda_{i|s_2}\right)}} + \frac{|\lambda_{j|s_1}-\lambda_{j|s_2}|}{\sqrt{\frac{1}{2}\left(\lambda_{j|s_1}+\lambda_{j|s_2}\right)}} \tag{6}$$

To simplify notation in the proof, we first introduce variables for the sum and difference responses, $\gamma_{i/j} = \lambda_{i/j|s_1}+\lambda_{i/j|s_2}$ and $\beta_{i/j} = \lambda_{i/j|s_1} - \lambda_{i/j|s_2}$, so that the inequality above becomes:

$$\frac{|\beta_i+\beta_j|}{\sqrt{\gamma_i+\gamma_j}} \leq \frac{|\beta_i|}{\sqrt{\gamma_i}} + \frac{|\beta_j|}{\sqrt{\gamma_j}}.$$

Multiplying by all denominators yields

$$|\beta_i+\beta_j|\sqrt{\gamma_i\gamma_j} \leq |\beta_i|\sqrt{\gamma_j(\gamma_i+\gamma_j)} + |\beta_j|\sqrt{\gamma_i(\gamma_i+\gamma_j)}.$$

Under the assumption that the two neurons in the multiunit have the same stimulus preference, $|\beta_i+\beta_j| = |\beta_i|+|\beta_j|$, and one can rearrange the terms as

$$|\beta_i|\left(\sqrt{\gamma_i\gamma_j+\gamma_j^2}-\sqrt{\gamma_i\gamma_j}\right)+|\beta_j|\left(\sqrt{\gamma_i\gamma_j+\gamma_i^2}-\sqrt{\gamma_i\gamma_j}\right) \geq 0.$$

Hence, we conclude that in the limit when the modulation is weak and neurons share the same stimulus preference (and thus same sign for optimal decoding weights) the informativeness of a multiunit is upper bounded by that of its component units. Using a similar derivation, one can also show that the informativeness of a multiunit is lower bounded by the average of the informativeness of the two neurons that compose it:

$$\frac{|\lambda_{i|s_1}+\lambda_{j|s_1}-\lambda_{i|s_2}-\lambda_{j|s_2}|}{\sqrt{\frac{1}{2}\left(\lambda_{i|s_1}+\lambda_{j|s_1}+\lambda_{i|s_2}+\lambda_{j|s_2}\right)}} \geq \frac{1}{2}\left(\frac{|\lambda_{i|s_1}-\lambda_{i|s_2}|}{\sqrt{\frac{1}{2}\left(\lambda_{i|s_1}+\lambda_{i|s_2}\right)}} + \frac{|\lambda_{j|s_1}-\lambda_{j|s_2}|}{\sqrt{\frac{1}{2}\left(\lambda_{j|s_1}+\lambda_{j|s_2}\right)}}\right) \tag{7}$$

While these constraints are derived under extreme assumptions, we can show numerically that the same intuition holds in the more relevant scenario when the modulation is not negligible (Fig. S3). For moderate modulation strength, $\sigma_m = 1$ and $c = [0,1]$, and random targeting ($c$ coupling assigned uniformly randomly to neurons with different tuning properties), the summed informativeness, $|d'_i + d'_j|$, lower bounds the multiunit informativeness for both pairs of neurons with aligned stimulus preference and pairs of neurons with dissimilar tuning (Fig. S3F). Note that tuning similarity, as measured by the inner product of the individual units' tuning functions, does not strongly affect multiunit informativeness (Fig. S3G).

Finally, we find the process of pooling neural responses alone cannot induce positive correlations between modulation and informativeness. The effect of modulation on a multiunit, as estimated via the modulator-guided decoder, takes the form $\mathbb{E}\left[(k_i+k_j)m\right] = \sigma_m^2\left(\lambda_i c_i \exp\frac{\sigma_m^2 c_i^2}{2} + \lambda_j c_j \exp\frac{\sigma_m^2 c_j^2}{2}\right)$. This estimated modulator coupling does not show targeting towards informative multiunits if the single unit coupling is unrelated to the single unit informativeness. When the modulator is targeted, some of this structure is preserved on the multiunit level (Fig. S3H). This is why using our modulator-guided decoder on the V1 data still performs well. Overall, these results suggest that analyzing multiunits underestimates the true informativeness of the underlying neurons. In itself it does not induce dependencies between informativeness and modulation, as used by the decoder proposed in the theory.

**Impact of multiunits on the estimation of modulator targeting.** Last, we wanted to understand how the presence of multiunits impacted the fitting and interpretation of our model. The following analysis was performed for simulations of either single or multiunits and the results regarding targeting structure were equivalent. Only the multiunit scenario is reported as its results include the single unit case.

To investigate the impact of multiunits on the model fitting procedure, we use the same modulated SR model that pools pairs of neurons, with various degrees of informativeness. We simulate a population of multi-units with data-matched statistics (in terms of number of units and firing rate distribution) including pairs of neurons that are either coupled to the global modulator or not (Fig. S3E). We consider two targeting scenarios: 1) preferential targeting towards task informative neurons, as hypothesized in the theory (Fig. S3I-L) and 2) random targeting, with coupling strengths that are independent of neuron informativeness (Fig. S3M-P). We apply the data analysis pipeline used on the experimental data to the resulting artificial datasets to fit a modulator and assess the properties of the corresponding PLDS estimated coupling strengths.

In the first scenario, we have a diversity of degrees of informativeness for the single units (Fig. S3I) and strong correlations between single unit modulation coupling and informativeness (Fig. S3J). The corresponding estimated multiunit informativeness remains upper bounded by the sum of the informativeness of the component neurons, as before. The correlation between modulator coupling and informativeness is weaker than for single units, but follows the same trend (Fig. S3K). This suggests that if targeting is presented in the single neurons, then analysis of multiunit measurements is likely to underestimate the degree of targeting.

The picture is quite different when no targeting is enforced (specifically, modulator couplings are random and independent). Somewhat counter-intuitively, since higher modulator coupling introduces noise and decreases a neuron's informativeness, random targeting leads to an anti-correlation between informativeness and coupling strength (Fig. S3N). The sum of single unit $d'$ values remains an upper bound for multiunit informativeness, with less variability than in the targeted scenario (Fig. S3O). Importantly, there are no spurious correlations in the estimated between multiunit coupling and informativeness when structured targeting is not present in the single units. Hence, the presence of multiunits alone cannot explain the modulator targeting we find in the data.

In the V1 data, different multiunits are likely to have different numbers of neurons driving their responses. This variability will further bias estimates of modulator strength towards larger multiunits. The opposite effect occurs with informativeness: larger multiunits will in general have more diverse responses, reducing their estimated informativeness. Thus, variability in multiunit size induces *negative* correlations between modulation strength and informativeness. In summary, it is likely that our empirical estimates based on the data underestimate the degree of targeting V1 neurons.

### S10 Theoretical results for modulator-guided decoding

**Encoding analysis** If the modulator coupling is unstructured (e.g., $c_n$ are distributed randomly), then the modulator can be viewed as a global noise source that decreases the discriminatory power of V1 responses proportional to its variance. If the modulator coupling is targeted towards informative neurons (e.g., $c_n$ proportional to difference in mean responses to stimuli), then the harmful effect becomes stronger, as noise is introduced specifically where it most impacts the encoded signal. We quantify the signal-to-noise ratio, using a Fisher Linear Discriminant:

$$\mathrm{SNR} = \frac{\left(\mathbf{a}^\top(\mu_\mathbf{1} - \mu_\mathbf{0})\right)^2}{\mathbf{a}^\top\Sigma_1\mathbf{a} + \mathbf{a}^\top\Sigma_0\mathbf{a}}, \tag{8}$$

where $\mathbf{a}$ denotes the decoding weights, and $\{\mu_s, \Sigma_s; s \in [0,1]\}$ are the population mean and covariance for the two stimuli. We evaluate this measure for the optimal decoding weights $\mathbf{a} = \mathbf{a}^{(\mathrm{MC})}$(see main text, optimal decoding weights, Eq. 2) and find that discriminability (SNR) decreases faster if modulation is targeted (Fig. S4A). A more detailed analysis of how encoding is affected by modulator strength and targeting can be found in [11]. Simulations were implemented in python and the code reproducing the figures is available on `https://github.com/CarolineHaimerl27/modulator_guided_decoding`.

**Fraction of informative neurons** Intuitively, the problem of finding the subset of task-informative neurons in a population becomes more difficult as the proportion of informative neurons decreases. To assess this, we simulated a population of neurons (Fig. S4B), varying the percentage of neurons that are task-informative. Mean firing rates of all neurons in the population were the same, but the firing rates of informative neurons were stimulus-modulated by $\pm 5\%$. We found that the modulator-guided decoder matched the performance upper-bound for the entire range (Fig. S4C). In contrast, the performance of the sign-only decoder is suboptimal, and suffers as the number of neurons that are uninformative increases (Fig. S4C, gray). This is expected given that the sign-only decoder groups all neurons into two subpopulations based on stimulus preference, and simply compares their total firing rates. When all neurons are similarly informative, the decoder performs optimally, but otherwise performance is well below the ideal observer bound. If we were to include inactive units (i.e. uninformative neurons with very low activity which correspond to neurons with receptive fields away from the task relevant stimuli) the overall performance of the decoders would decrease – as inactive units add noise– but it would not change these results qualitatively (for details see [11]).

**Learning the signs of decoding weights** We have separated the problem of approximating the optimal decoding weights into two sub-problems: estimating the magnitudes of the weights, $|a_n|$, and estimating their corresponding signs (i.e. their preferred stimulus). For the modulator-guided decoder, the first estimation happens within trials, based on correlations between individual neural responses and the modulator, whereas the signs are learned from explicit feedback given at the end of each trial. Here we simulate informative neurons with different strengths of modulation and examine the correctness of sign estimation as a function of number of training examples (Fig. S4D). We find that, in general, the number of trials needed to estimate the signs is small, about 10 trials for the moderate modulator strengths in our simulations. Hence, if the decoding mechanism can identify the few informative neurons and attribute negligible weights to the rest, finding the signs of the informative neurons is fast.

**Size of population** We varied the size of the simulated population while keeping the % of informative neurons fixed at 5%. We find that MG decoding qualitatively performs similar when population size is increased to $N = 2000$, $N = 4000$ and $N = 10000$ neurons (Fig. S4E-G).

### S11 Learning MG weights and signs jointly via eligibility traces

We illustrate a potential learning rule for estimating the modulator-guided decoding weights in a biologically plausible way, using eligibility traces online learning. The key idea is to use eligibility traces, updated online on the time scale of modulator fluctuations, to estimate the degree of modulation of individual neurons, then combine these correlations with the explicit task feedback received at the end of each trial.

First, one eligibility trace integrates evidence of modulation in neuron $n$ over time $t$ which is independent of the stimulus presentation.

$$e_{n,t+1} = \alpha e_{n,t} + (1 - \alpha) k_{n,t} m_t. \tag{9}$$

On the same time scale, we use another eligibility trace for the overall firing rate of neurons to correct for the bias (see in Methods of main text)

$$r_{n,t+1} = \alpha r_{n,t} + (1 - \alpha) k_{n,t} \tag{10}$$

Second, information about the signs of the decoding weights comes from trial feedback and is tracked down in the form of a rescaled error:

$$b_{n,i} = (\hat{s}_i - s_i) \cdot \hat{k}_{n,i}, \tag{11}$$

where $\hat{k}_{n,i}$ denotes the total spike count during the $i$th stimulus presentation, and $\hat{s}_i = \sum_n \hat{w}_n \hat{k}_{n,i}$ is the estimated stimulus category.

Whenever an error occurs in trial $i$ (decision made about a stimulus was incorrect) we make an error-dependent update; if no negative task-feedback is provided, only the amplitude of the MG weight is updated, while the sign is preserved.

$$\Delta c_n = b_{i,n} \frac{e_{n,t}}{r_{n,t}} + \delta(b_{n,i}) \mathrm{sign}(\hat{w}_n) \frac{e_{n,t}}{r_{n,t}} \tag{12}$$

where $\delta$ is the Kronecker delta. $\frac{e_{n,t}}{r_{n,t}}$ corresponds to the estimation of the absolute MG weights and is combined with $b_{i,n}$ which corresponds to the gradient of a regression loss. Finally, these changes are integrated to provide the estimated decoding weight:

$$\hat{w}_n = \gamma \hat{w}_n + (1 - \gamma) \Delta \hat{w}_{n,i,t} \tag{13}$$

In numerical simulations, we find that this learning rules allows a reasonably robust online estimation of decoding weights (Fig. S4H). Specifically, we simulated a population of 100 neurons for 100 stimulus presentation, each lasting 200ms, at a time resolution of 50ms. Neural responses during a stimulus presentation are generated according to our encoding model:

$$k_{n,t} \sim Poisson\left(\exp(\lambda_n(s)) \exp(c_n m_t)\right) \tag{14}$$

with the modulator drawn independently from a zero mean, unit variance Gaussian distribution. The stimulus-response rate $\lambda_n(s)$ is set so that neurons differentiate between the two task categories to varying degrees, with about half the population preferring one stimulus and the other half the other stimulus (Fig. S4I). The eligibility traces that contribute to the estimation of the absolute weight of $\hat{w}_n$, Eq. 9 and Eq. 10, integrate information with a time constant given by $\alpha = 0.9$, while the weight updates happen at the time scale of stimulus presentations, with a learning rate $\gamma = 0.999$. Estimated weights $\hat{w}_n$ converge to values that preserve the ground truth after only 20 stimulus presentations (Fig. S4I), with decoding performance showing around 90% accuracy.

### S12 Details on comparison of modulator strength

We find that the overall modulator strength relative to the stimulus drive decreases in the control condition compared to the relevant tasks.

This difference is driven by a change in the modulator strength, which decreases in the control (non-parametric Wilcoxon U-test $p \ll 0.0001$). The stimulus (contrast) induced variance (variance in stimulus ) is relatively constant across the task-conditions ($p > 0.05$) (Fig. S5A-B).

### S13 Relationship of other noise sources to behavior

We have found that the modulator coupling is higher in neurons that also show a strong correlation with the behavioral choice of the monkey, indicating that they are preferentially recruited for the decision. As a control, we want to know whether this relationship extends to other shared sources of noise in the population. We take advantage of the fact that the first PC is correlated with the PLDS extracted modulator, while the second PC is uncorrelated with the modulator and accounts for a very similar fraction of variance explained. We repeat the original analysis with PC1 as a proxy for the modulator and PC2 as another shared noise sources. As already reported for the modulator, PC1 is predictive of a unit's correlations to behavior, but this effect does not extend to the second PC (Fig. S5C). This dissociation supports the idea that the relationship between single cell fluctuations and behavior is specific to the modulator and not to other sources of noise in the neural responses.

### S14 Cross-task decoding

Here we test whether optimal decoding weights computed on the data from one relevant task condition block in a session can also decode in another block of the same session. Specifically, we are interested in whether the decoding weights computed in one relevant task can also be used to decode in the other relevant task. Fig. S5D shows that decoding from a different block with a different task condition does not perform much above chance, while decoding from a different block with the same task condition performs significantly better (t-test, p<0.0001). This suggests that the optimal decoding weights for one relevant task are substantially different from the ones for the other relevant task.

### S15 Extension on MT single unit analysis

Some recordings include a single unit in area MT. These units vary in their informativeness for the task (Fig. S6A). The stimulus response of MT units is modeled by the same SR model as used for V1, but augmented to include the drift direction of the stimulus. We verify the model fit over multiple cross-folds and find almost exclusively good model fits quantified by a comparison to a constant rate model through the pseudoR measure (Fig. S6B).

While most V1 modulators extracted from the data show a strong targeting towards informative neurons, a few outliers do not. We look for differences between targeted and untargeted modulators with respect to their predictability for MT. We find that only those V1 modulators that are well targeted to informative neurons have predictive power for their respective MT units (Fig. S6C). This could be because of differences in fit qualities where for some blocks the estimated V1 modulator coupling is too noisy and hence does not reveal targeting and also prevents modulator estimates precise enough to be predictive of MT. Alternatively, the untargeted V1 modulators may reflect a different kind of shared noise in the population that is private and not propagated to MT.

### S16 Extension on MT population analysis

The modulated SR model is fit to a population of 24 units. The SR model includes direction and contrast. The fit of the SR model is good across all blocks compared to a constant rate model (Fig. S6D). The modulator further improves this fit in all but one block (Fig. S6E). The modulator is not significantly dependent on the stimulus contrast in 72% of blocks (Fig. S6F). For the other 28% of blocks the modulated SR model does not manage to separate stimulus response and the modulator. We exclude those blocks from the analysis to avoid confounds.

### S17 Supplementary Figures

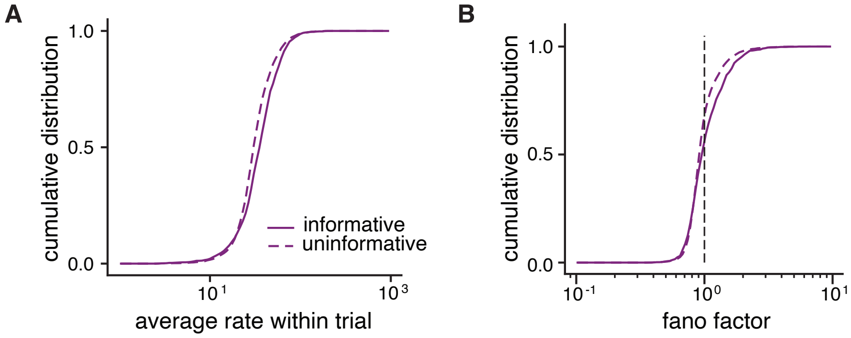


Figure S1: Basic statistics of V1 units. A) Average firing rate during a trial, separating the units into informative and uninformative subpopulations. B) Fano factor distributions for informative and uninformative units.

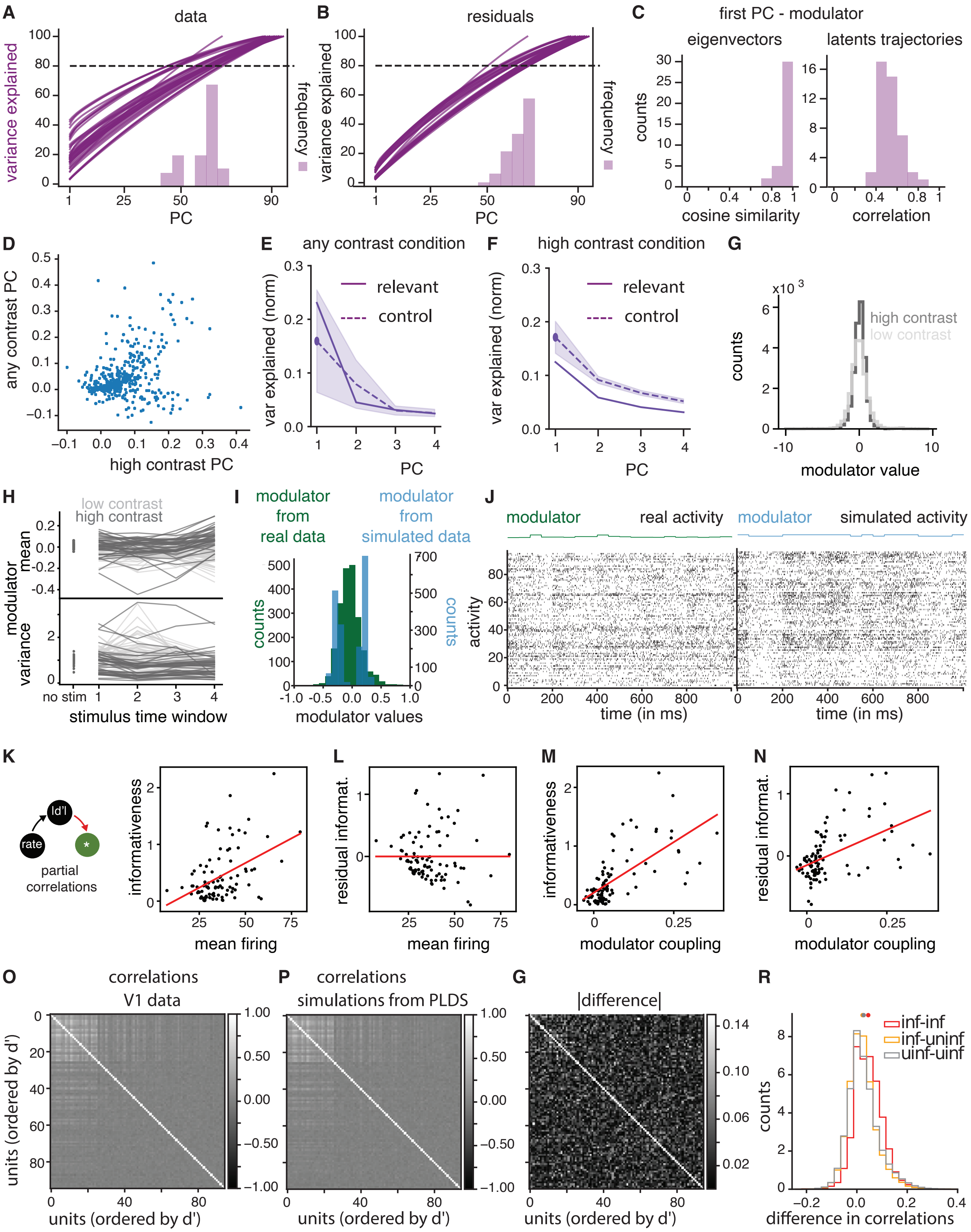


Figure S2: (Caption next page.)

Figure S2: A-F) PCA analysis of V1 population activity. Variance explained by principle components of A) population responses and B) SR residuals; we apply PCA to the concatenated trials of a block, each datapoint is the activity of a unit in a 50ms time bin. Histograms show the minimum number of PCs required to explain 80% of the variance in A) the data and in B) the SR residuals. C) Left, we take the cosine similarity between the eigenvector corresponding to the first PC of the residuals and the modulator coupling for every block. Here we plot the distribution over all blocks. Right, we compute the correlation between the trial-concatenated modulator found by the PLDS and by PCA on the residuals. We plot the distribution of correlation coefficients over all blocks. D) First PC computed on all stimulus presentations in the control task, over first PC computed on high contrast stimulus presentations. Each point represents a unit's loading on the PC axis. The graph pools over all control task blocks. If a population's first PC had a negative mean (mean of first eigenvector $< 0$) the entire vector is rotated by $-1$ to increase visual comparability. E) Normalized variance explained for first 4 PC axis extracted from residuals of any stimulus presentations using the SR model fitted, for either the relevant or the control tasks. Lines show averages over all blocks and shaded region shows the sd. F) As H but for high contrast stimulus presentations. G-H) Inferred modulator statistics. G) The distribution of modulator values during high/low contrast stimulus presentations. H) Every line represents the mean (top) and variance (bottom) of the modulator in a block, estimated from the different time bins of a stimulus presentation and for low and high contrast. Dark grey lines represent high contrast and light grey low contrast presentations. I-J) On/Off states I) Modulator distribution extracted from an example block population (green) or extracted from simulations from the same model fit but using an artificial bimodal modulator instead (blue). J) Population spiking activity for for one second of that same example block (left) and simulated version (right). K-N) Partial correlation analysis for mean rate, coupling and informativeness. K) Dependence between |d'| and mean firing. L) Residuals of linear fit as a function of firing rate are unstructured. M) The relationship between informativeness and coupling. N) The same for residual informativeness (unexplained by differences in mean firing). O-R) Excess noise correlations. O) Pairwise noise correlations of a population in an example block computed on high contrast stimulus presentations. The color bar indicates Pearson correlation coefficient. P) Pairwise noise correlations in simulations from the same example modulated SR model. G) Difference between pairwise noise correlations in data (A) and in simulations (B). Colors indicate difference in Pearson correlation coefficient. R) Distribution of differences in pairwise correlation coefficients over all blocks. Colors indicate the type of pairs; Red for two informative units, yellow for one informative and one uninformative units, grey for two uninformative units.

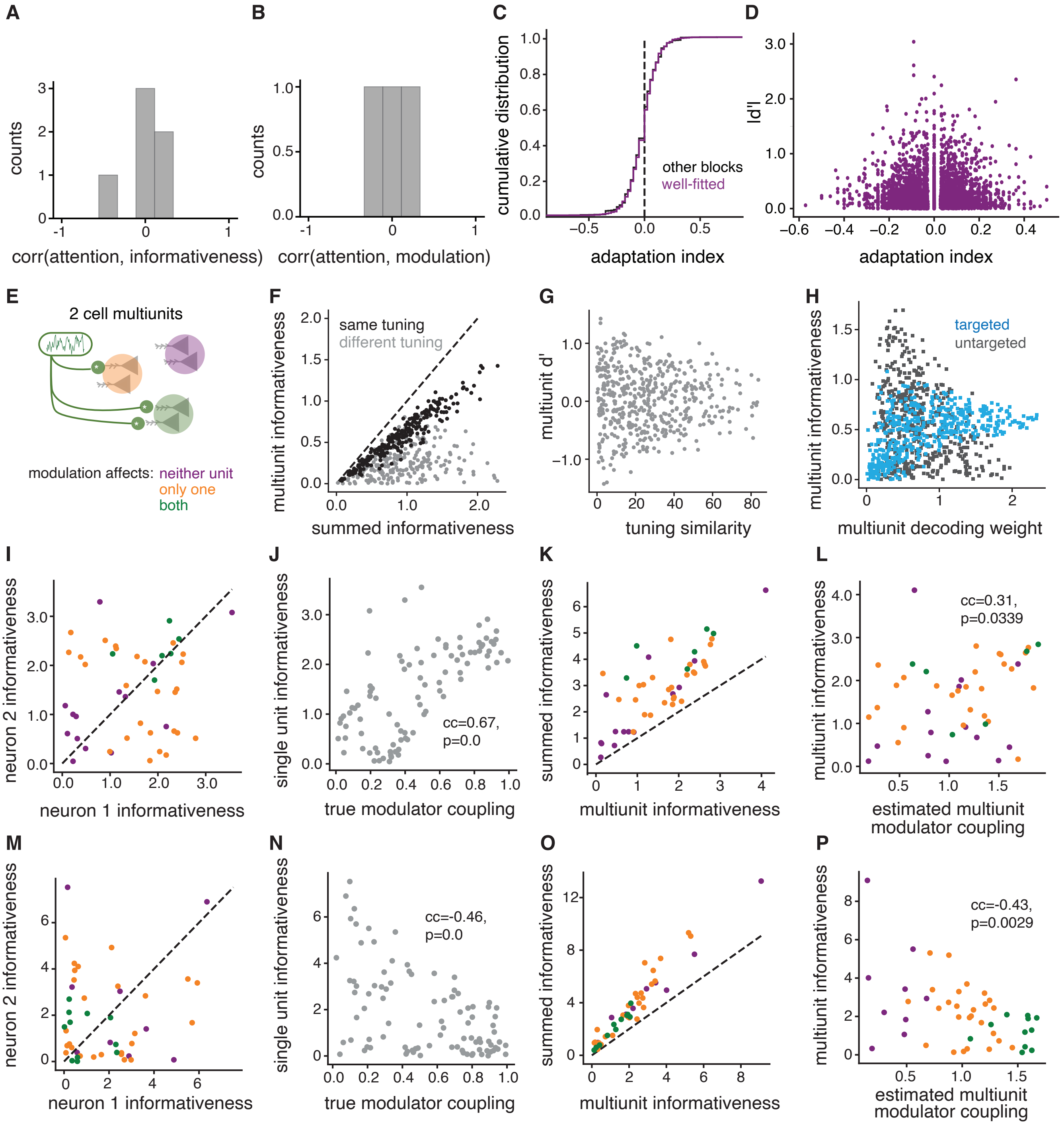


Figure S3: (Caption next page.)

Figure S3: A-B) Distribution of Spearman correlation coefficients over blocks between A) attentional modulation index and informativeness and B) between attentional modulation index and modulator coupling. C-D) Effects of adaptation. C) The distribution of adaptation indices for blocks well fit by mod-SR (purple) and all blocks (black). D) Informativeness of each unit, measured by $|d'|$ as a function of the unit's adaptation index; no linear dependency, Spearman $p = .36$. E-M) Effect of multiunits on key analysis measurements. E) We model multiunits by summing over the activity of two model Poisson neurons with modulated rates. F) Informativeness of multiunit $|d'_{(i,j)}|$ versus the sum of informativeness of the component units, $|d'_i| + |d'_j|$. G) Multiunit $d'$ as a function of the cosine similarity between the tuning of the individual neurons. H) Multiunit informativeness as a function of estimated multiunit modulator-guided decoding weights for a simulated targeted and untargeted population. I) The informativeness of individual units comprising the set of simulated multiunits; colors mark type of modulation for each pair, as in A). J) Corresponding modulator couplings are correlated with single unit informativeness (the 'targeted modulation' scenario). K) Multiunit informativeness versus sum of single neurons $|d'|$. L) Modulator coupling estimated using the PLDS model and its correlation to multiunit informativeness. M) to P) as above, but for the 'untargeted' scenario.

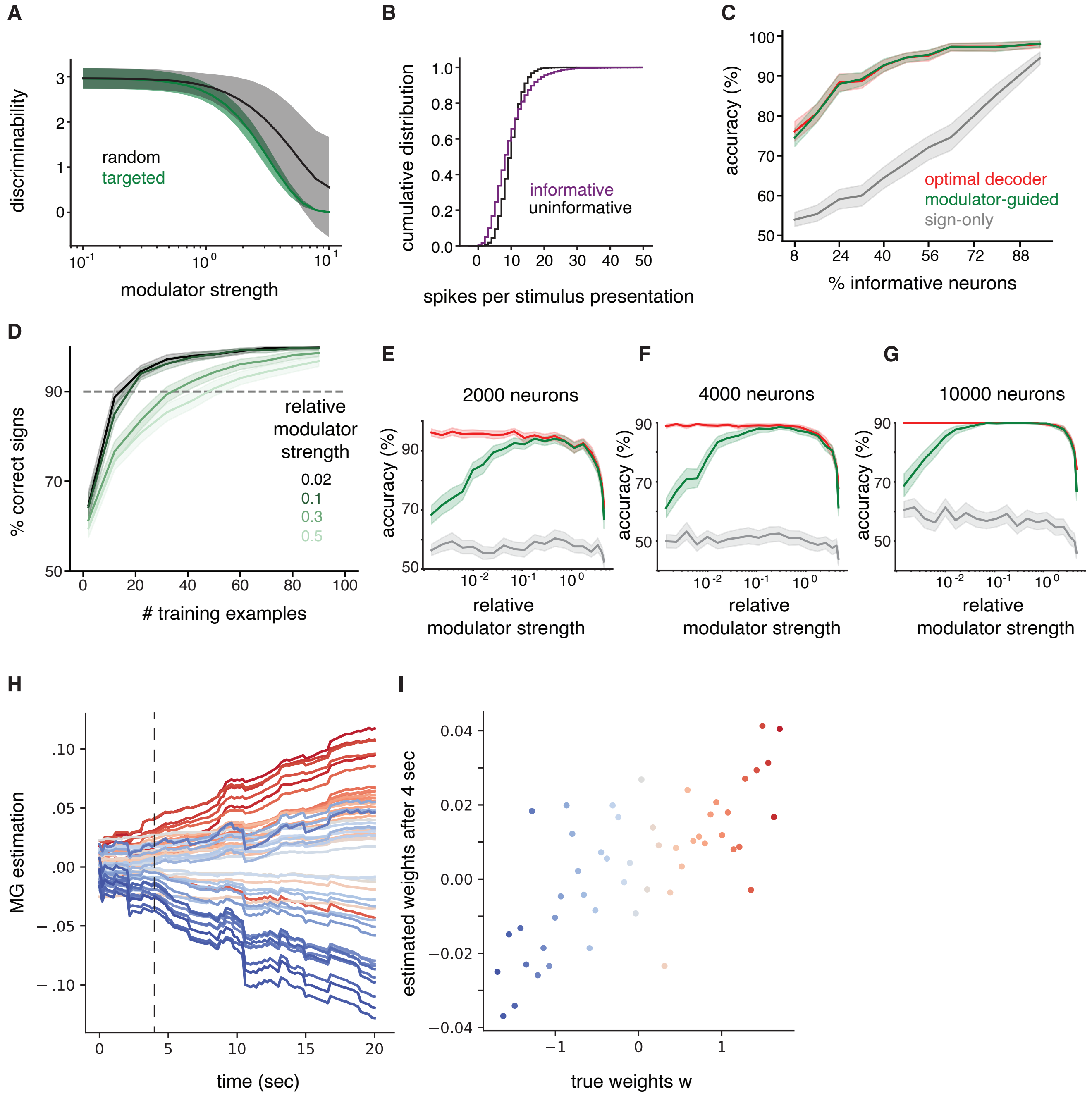


Figure S4: A-E) Simulations of decoding from V1. A)The effect of modulation on stimulus SNR, as measured by the Fisher Linear Discriminant, for unstructured and targeted modulator coupling. N=100 neurons, 50 inactive, 12 informative, 38 uninformative. B-C) Effect of fraction of informative neurons on decoding performance. B) Firing rate distributions in a simulated population; all neurons are similarly active, but uninformative neurons do not change their responses as a function of the task relevant stimuli while informative neurons are modulated by ±5%; N=50 neurons. C) Decoder performance as a function of the fraction of informative neurons (constant total population of 50 neurons, for details see text). D) The percentage of correctly estimated decoding signs as a function of the number of training examples. Different colors correspond to varying relative modulator strengths (see Methods for details). E-G) Performance with varying size of the population. E) As main Fig. 3D but using 2000 instead of 1000 neurons. F) and G) as A but with increasing population size. H-I) Eligibility Trace H) Estimation of weights $\hat{w}_n$ over learning; each stimulus presentation lasts 200ms. Individual lines correspond to decoding weights of 100 neuron. Color gradient indicates the rank of the corresponding ground truth decoding weight in the population, with red and blue representing opposite tuning preferences. Dashed line indicates minimal training of 4 seconds, or 20 stimulus presentations. I) Final estimates $\hat{w}_n$ after 4 seconds of learning compared to the optimal decoding weights. Colors as in A. Both axes are z-scored.

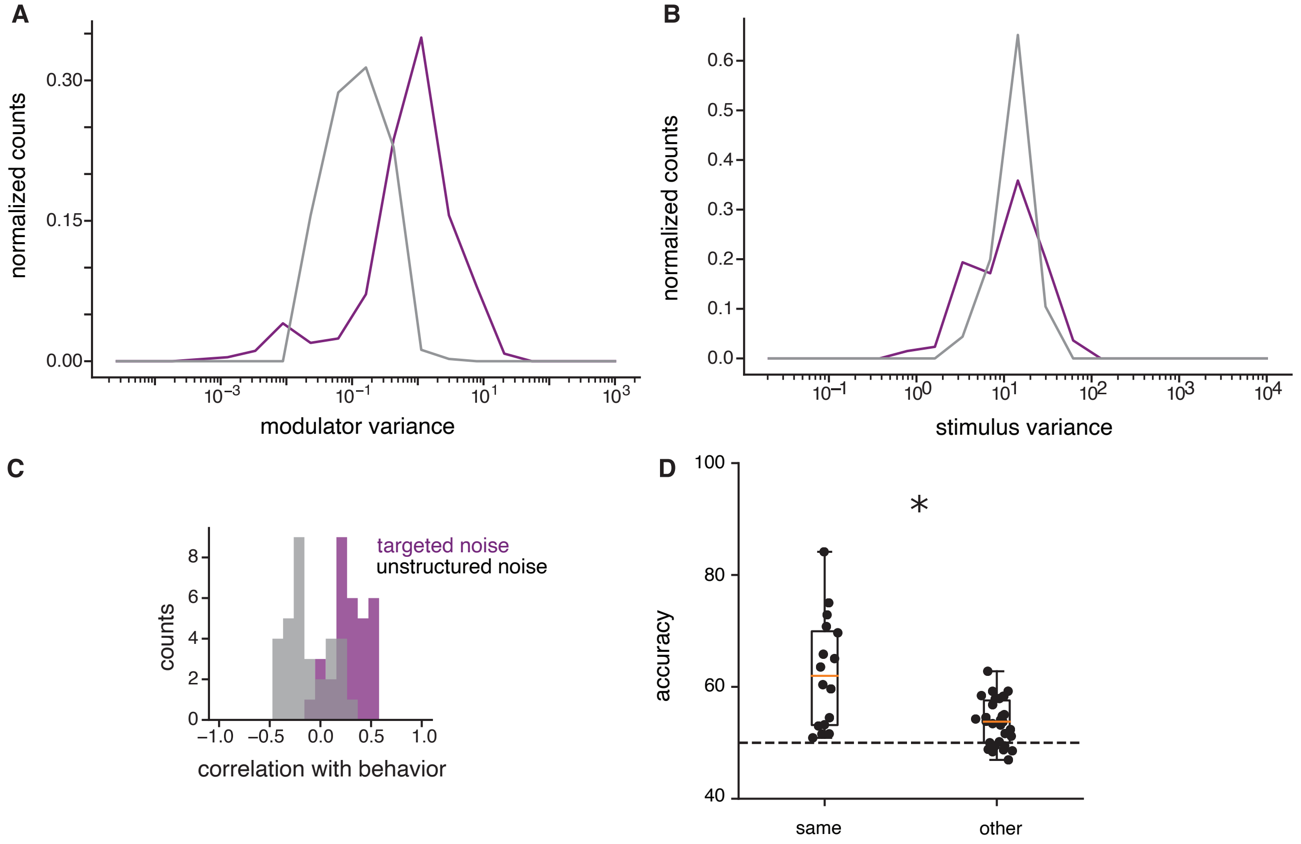


Figure S5: A-B) Modulator strength and stimulus strength analyzed separated. A) Modulator strength (variance of modulator with unit vector coupling) in relevant (purple) versus control (grey) task over all blocks. B) Stimulus induced variations in relevant and control task. D) Cross-task decoding. Here we compute the optimal decoding weights for one block of a relevant task condition and apply it to another block in the same session, which can be either of the same task condition (same) or of the other relevant task condition (other). C) Relationship of other noise sources with behavioral correlation. We plot the distribution of correlation coefficients across blocks between behavior and the first PC (purple) or the second PC (grey).

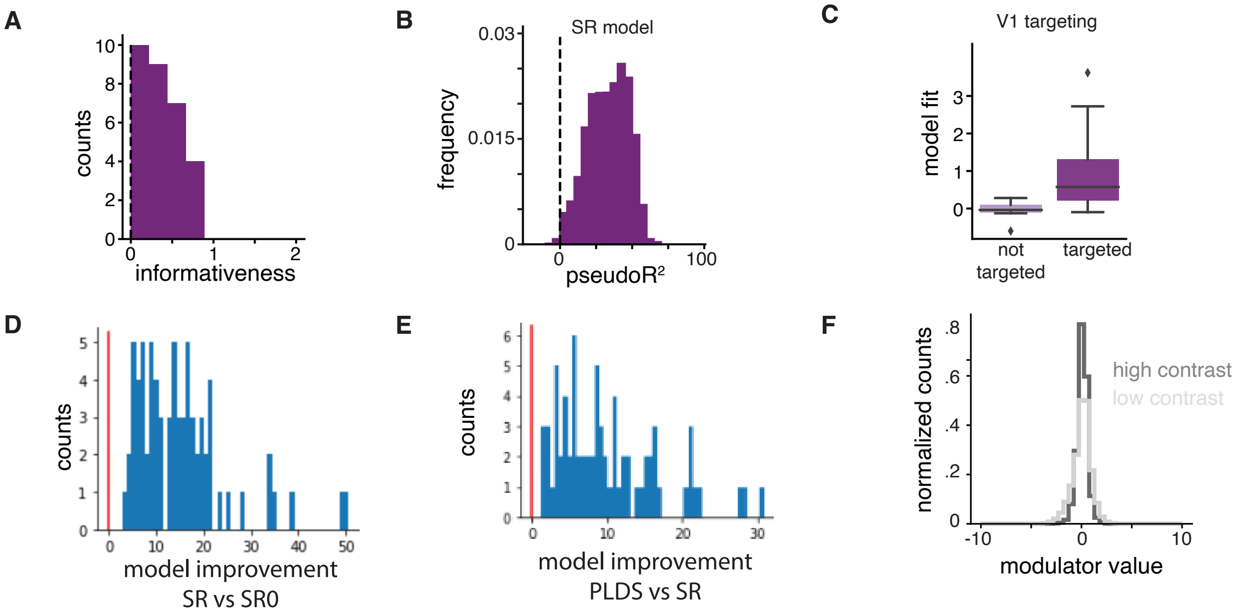


Figure S6: A) Distribution of informativeness ($|d'|$) over all MT units from the single unit recordings. B) The fit quality for the MT SR model is quantified by pseudoR which compares the log-likelihood of the SR model against a simpler constant rate Poisson model. We plot the distribution of pseudoR values for all units and different cross-folds. C) Blocks are split into subsets for which the estimated V1 modulator targets preferentially informative neurons (as measured by significant correlations between modulator coupling and informativeness) and blocks without significant targeting. We plot the respective distributions of model fit quality (pseudoR). D-F) Model fit for population MT recordings. D) Average log-likelihood fit for the SR model for each block population compared to a constant rate model. E) Average log-likelihood fit for the modulated SR model for each block population compared to the SR model. F) The distribution of MT modulator values during high/low contrast stimulus presentations.

### References


[1] Yu, B. M. *et al.* Gaussian-Process Factor Analysis for Low-Dimensional Single-Trial Analysis of Neural Population Activity. *Journal of Neurophysiology* **102**, 614–635 (2009).

[2] Engel, T. A. *et al.* Selective modulation of cortical state during spatial attention. *Science* **354** (2016).

[3] Kanitscheider, I., Coen-Cagli, R. & Pouget, A. Origin of information-limiting noise correlations. *Proceedings of the National Academy of Sciences of the United States of America* **112**, E6973–82 (2015).

[4] Moreno-Bote, R. *et al.* Information-limiting correlations. *Nature Neuroscience* **17**, 1410–1417 (2014). `arXiv: 1411.3159v1`.

[5] Treue, S. & Martínez Trujillo, J. C. Feature-based attention influences motion processing gain in macaque visual cortex. *Nature* **399**, 575–579 (1999).

[6] Maunsell, J. H. & Cook, E. P. The role of attention in visual processing. *Philos Trans R Soc Lond B Biol Sci.* **357**, 1063–72 (2002).

[7] Ruff, D. A. & Cohen, M. R. Attention can either increase or decrease spike count correlations in visual cortex. *Nature neuroscience* **17**, 1591–7 (2014).

[8] Ruff, D. A. & Cohen, M. R. Attention increases spike count correlations between visual cortical areas. *The Journal of neuroscience : the official journal of the Society for Neuroscience* **36**, 7523–34 (2016).

[9] Cohen, M. R. & Maunsell, J. H. Attention improves performance primarily by reducing interneuronal correlations. *Nature Neuroscience* **12**, 1594 (2009).

[10] Snyder, D. L. & Miller, M. I. *Random point processes in time and space* (Springer Science & Business Media, 2012).

[11] Haimerl, C., Savin, C. & Simoncelli, E. Flexible information routing in neural populations through stochastic comodulation. *Advances in Neural Information Processing Systems* **32**, 14402–14411 (2019).
